# Repository of logically consistent real-world Boolean network models

**DOI:** 10.1101/2023.06.12.544361

**Authors:** Samuel Pastva, David Šafránek, Nikola Beneš, Luboš Brim, Thomas Henzinger

## Abstract

Recent developments in both computational analysis and data-driven synthesis enable a new era of automated reasoning with logical models (Boolean networks in particular) in systems biology. However, these advancements also motivate an increased focus on quality control and performance comparisons between tools.

At the moment, to illustrate real-world applicability, authors typically test their approaches on small sets of manually curated models that are inherently limited in scope. This further complicates reuse and comparisons, because benchmark models often contain ad hoc modifications or are outright not available.

In this paper, we describe a new, comprehensive, open source dataset of 210+ Boolean network models compiled from available databases and a literature survey. The models are available in a wide range of formats. Furthermore, the dataset is accompanied by a validation pipeline that ensures the integrity and logical consistency of each model. Using this pipeline, we identified and repaired 400+ potential problems in a number of widely used models.

## Background & Summary

Logical models in general, and Boolean networks in particular, represent one of the fundament modelling frameworks employed by systems biology^1^. Historically, much of the Boolean network design and analysis has been performed either manually, or through computer assisted approaches requiring extensive involvement of domain experts^2–4^. However, recent developments in topics like attractor^5–7^ and trap space^8^ detection, bifurcation analysis^9^, and synthesis from data and prior knowledge^10,11^ enable comprehensive automated analysis of large, possibly partially unknown networks comprising hundreds of components. This proliferation of Boolean network tools and methods motivates the need for extensive and reproducible validation using realistic real-world models. Typically, this role is served by ad hoc subsets of models available in popular systems biology databases such as CellCollective^12^, Biomodels^13^ or GINsim^14^. However, this approach suffers from many obvious downsides.

While these databases have non-trivial overlap, they are mostly focused on specific human-curated models. As such, they are inherently limited in scope to the models recognised by the database curators. Furthermore, models available from multiple sources often contain subtle but critical modifications in the network structure and initial conditions. Finally, these sources often lack automated validation of the model integrity and logical consistency, meaning errors can propagate to models derived in the future, or go unnoticed when performing analysis.

To address these issues, we present the *Biodivine Boolean Models* (BBM) dataset. BBM covers 210+ models from an extensive set of sources, including the aforementioned databases and independently published models (see Figure 1). For each model, BBM tracks the primary source of the model (i.e. its publication and authors), any secondary sources (e.g. databases) and the source for the actual model files (e.g. git repositories, supplementary materials, database entries, etc.).

**Figure 1.**
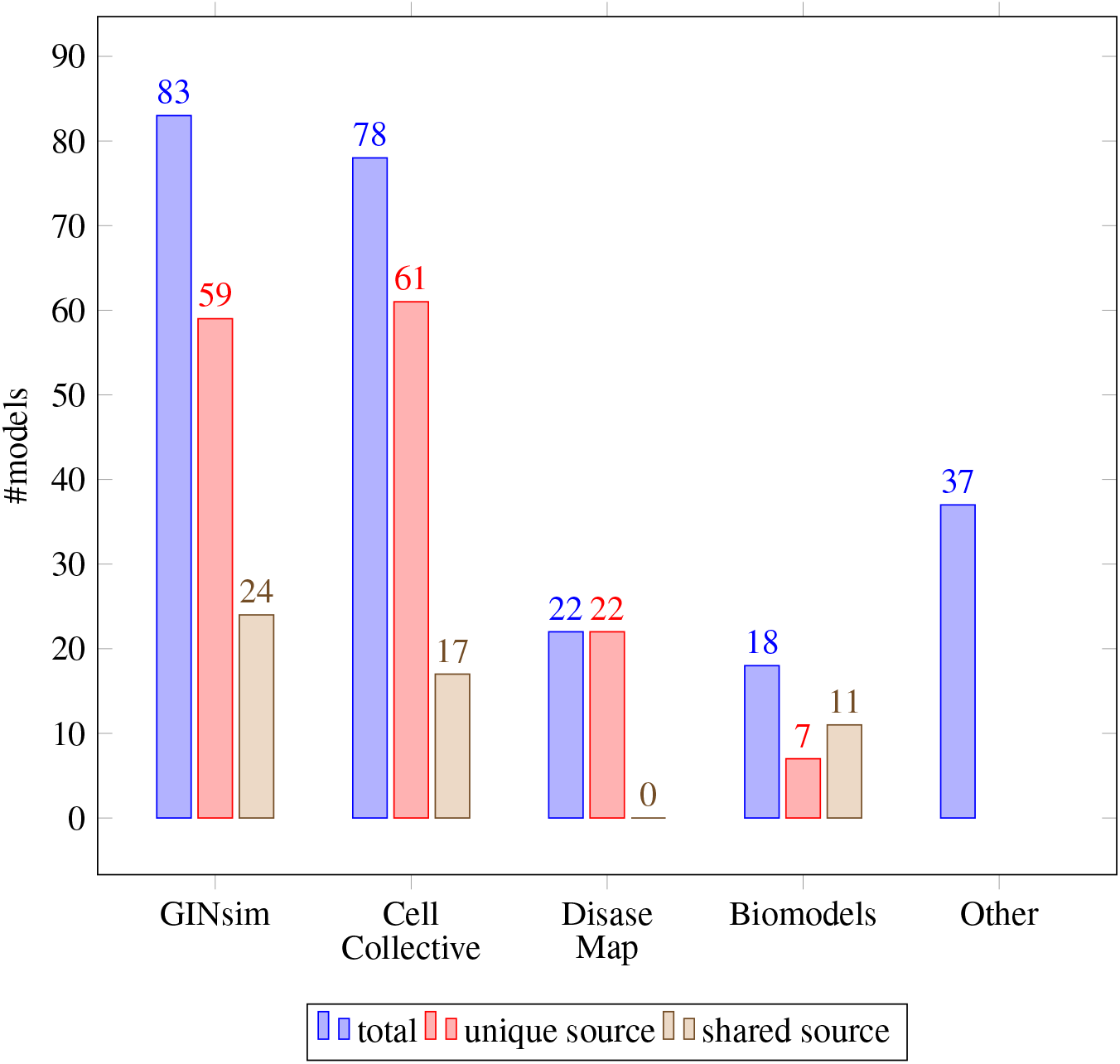
Models of the BBM dataset grouped by the database in which they appear, including the number of models uniquely available in each database and the models shared between multiple sources (see Online-only Table 2 for exact mapping).

Aside from model aggregation, BBM provides several benefits to tool authors: First, models tracked in BBM are available in several widely recognised formats (`bnet`^15^, `aeon`^16^, and `sbml`^17^) and can be downloaded in bulk. Second, BBM eliminates certain ambiguities in model representation, such as the treatment of network input nodes. Third, BBM performs extensive static validation to ensure logical consistency between variable update rules and the associated influence graph of each model. This process identifies and repairs common problems in logical models such as mismatch in monotonicity of the regulation and the update function, or the presence of (logically) redundant model components. Finally, BBM provides an automated reproducible workflow to generate custom variants of the dataset based on the user provided criteria.

Overall, we believe BBM significantly simplifies the process of validating and benchmarking Boolean network tools as it provides a large, clearly defined and validated set of models. To the best of our knowledge, BBM is also the largest publicly available set of real-world (i.e. not randomly generated) Boolean network models.

## Methods

In this section, we first give a quick overview of the relevant elements of Boolean network models. We then proceed to list how the actual BBM dataset is created, curated, validated and distributed.

### Boolean networks

First, let us give a brief formal introduction into the topic of Boolean network modelling. By 𝔹, we denote the set {0, 1}. We then say that a *Boolean network ℬ* over *n* variables is an indexed collection *ℬ* = {*F*_1_, …, *F*_*n*_}, such that each *F*_*i*_ is a Boolean *update function F*_*i*_ : 𝔹^*n*^ *→* 𝔹 (the set of Boolean vectors 𝔹^*n*^ is called the *state space* of *ℬ*). Naturally, within a real-world model, variables are also augmented with human-readable names and other biologically relevant metadata, while functions are described using Boolean expressions or function tables.

### Signed Influence graph

Individual update functions typically depend only on a small subset of the network variables, not the whole network state. This fact is communicated using the *signed influence graph* (often also called the *regulatory network*) associated with the Boolean network *ℬ*. This is a labelled directed graph *ℐ* = (*V, E, l*) such that its vertices are the variables of *ℬ, V* = {1,…, *n*}, edges *E ⊆ V ×V* represent dependencies between variables (we say that *x regulates y* when (*x, y*) ∈ *E*) and *l* : *E* → {⊕, ⊖, ?} labels each edge with an optional *monotonicity* (positive, negative, or unknown). In rare cases, *mixed* monotonicity (combination of ⊕ and ⊖) is used to signify that the relationship is expected to be non-monotonic. However, for simplicity and due to its rareness, we do not consider such case here and treat such *mixed* labels simply as *unknown*. The influence graph is typically provided directly by model authors.

### Properties of Boolean functions

Intuitively, the influence graph *ℐ* imposes additional requirements on the structure of the network’s update functions *F*_*i*_. Formally, we can understand these requirements as *essentiality* and *monotonicity* of individual function inputs. First, given a state *x* ∈ 𝔹^*n*^, we write *x*[*i* ← *b*] to denote a copy of *x* with the value of the *i*-th variable substituted for *b ∈* 𝔹. Then, given a function *F* : 𝔹^*n*^ *→* 𝔹, we say that the *i*-th variable is an *essential* input of *F* if the output of *F* depends on this variable, formally ∃*x* ∈ 𝔹^*n*^. *F*(*x*[*i ←* 0]) ≠ *F*(*x*[*i* ← 1]). We also say that *F* is *positively monotonic* in its *i*-th input when increasing this input cannot decrease the output of *F*, formally ∀*x* ∈ 𝔹. *F*(*x*[*i* ← 0]) ≤ *F*(*x*[*i ←* 1]). Symmetrically, we say that the input is *negatively monotonic* when ∀*x* ∈ 𝔹. *F*(*x*[*i* ← 0]) ≥ *F*(*x*[*i* ← 1]).

In practice, these essentiality and monotonicity requirements map to the biological assumptions about the modelled system. For example, if a transcription factor is observed to activate the expression of a particular gene in a wet lab experiment, we expect it to act as an essential and positively monotonic input in the corresponding update function of our in silico model. Overall, we expect that the essential inputs of each update function are exactly the regulators declared within the influence graph and that the monotonicity of these inputs matches the associated edge labels *l*.

## Model Acquisition

Next, we describe the process of compiling the BBM dataset. Due to the high variability of model sources, the process of incorporating a new model is largely manual (aside from validation and consistency checking covered in the next section).

### Eligible models

To admit the widest possible assortment of realistic models, we currently allow submission of any logical model that can be reliably retrieved from a known database, repository, or an associated publication. The only requirement is that the model must be based on a real biological system. That is, we do not accept randomly generated or otherwise synthetic models. However, we do not require any specific level of curation or validation of any biological claims. In particular, the model can be hand made, automatically synthesised, or anything in between. Similarly, we accept many different model formats and we are willing to assist with translation if a machine-friendly model representation is not available.

### Multi-valued networks

The dataset also incorporates multi-valued networks. That is, logical models admitting multi-valued or bounded integer variables (as opposed to just Booleans). To maximise compatibility with the available tools, every such model is instead provided in a Booleanised representation. That is, the model is transformed into a plain Boolean network such that each multi-valued component with domain {0,…, *k*} is expanded into *k* Boolean components using the well-known Van Ham encoding^18^. This process is facilitated by the tool `biolqm`.^19^ Each Booleanised multi-valued model is distinguished using a `multi-valued` keyword included in the model’s metadata. Whenever possible, we also incorporate the `sbml`^17^ representation of the original (i.e. non-Booleanised) multi-valued model. However, such original files currently cannot be processed by the automated validation pipeline.

### Sources

As of 2022, the dataset incorporates 212 models from a wide range of publications. Notable sources include GINsim^14^ (83 models), CellCollective^12^ (78 models), Biomodels^13^ (18 models) and the COVID-19 Disease Map^20^ (22 models). If a model appears in multiple databases, the dataset lists such model only once with multiple source links, even if there are minor differences between the model files in each database. The contribution of each database to the overall dataset is visualised in Figure 1. Furthermore, Online-only Table 1 gives the full list of models including references to their original publications and Online-only Table 2 maps relevant database entries to the corresponding BBM models. The remaining models are sourced through a comprehensive literature search for Boolean network related keywords across PubMed and Google Scholar.

## Normalisation, Validation and Repair

Within BBM, each model is initially assigned a unique integer identifier (ID) and a human-readable name. Additionally, an *input* model file is required, using one of the eligible formats: `bnet`^15^, `aeon`^16^, and `sbml`^17^. In some cases, such representation is not available. For example, some models are described directly within the text of a supplementary material of a publication. For such models, we perform a manual conversion into one of the supported formats, but we also maintain the original model representation for future reference.

Our automated pipeline then translates the input model file into all three formats and performs several normalisation steps to maximise compatibility with other tools. First, we ensure that all variable names are unique and do not contain any special characters (in particular for `sbml`, model variables are further distinguished by separate IDs, hence a name can be an arbitrary string). Then, several normalisation and validation steps are performed to ensure the model contains no additional issues.

### Input node normalisation

A variable of a Boolean network is typically called an *input* if it has no incoming regulations (no incoming edges within the influence graph). Consequently, update function of such variable can be only *true* or *false*, since it cannot have any essential inputs. The particular choice of update functions then constitutes the *initial conditions* of the model. However, some authors also consider inputs to be constant but unknown, meaning their update function is the identity function. In such case, the variable has a positive self-regulation, but this is sometimes omitted if clear from context. The initial conditions of the model (if any) then need to be provided separately. Finally, some tools also allow to omit the update function of input nodes entirely, giving yet another possible representation for constant but unknown inputs.

The distinction between these representations is subtle but crucial in many instances: For example, when considering attractor detection, a constant *true* or *false* restricts the search to the specified value. Meanwhile, if the input is constant but unknown, the result covers attractors for every possible combination of input values. The number of attractors thus increases exponentially with the number of inputs in such a scenario.

In our pipeline, we detect the presence of network input nodes and normalise them during export. By default, models are distributed without the update functions on all recognised input nodes. However, using the custom dataset export procedure (discussed in the later *Data Distribution* section), users can chose what representation of inputs best suits their needs (i.e. a constant value, identity update function, or no update function at all).

### Regulation monotonicity and essentiality

Another step in our pipeline is responsible for validating the declared influence graph of the network against the actual update functions employed by the model. We do this using the symbolic BDD framework within the tool `aeon.py`^21^. This procedure allows us to formally verify the update function properties as described in the *Boolean networks* section. If an inconsistency is found, we manually adjust the influence graph such that the problem is removed, but we note this change in the metadata of the model. As such, the actual Boolean dynamics of the model are unchanged, but the error is recorded for future reference. The following errors can be detected and repaired using this procedure:

1. Input *j* is essential in *F*_*i*_, but there is no edge (*j, i*) *∈ E* in the influence graph; repaired by adding this regulation edge.
2. Regulation (*j, i*) *∈ E* appears in the influence graph, but *j* is not an essential input of *F*_*j*_; repaired either by removing the regulation edge, or by annotating the regulation as *non-essential* (only some output formats support such annotation).
3. Regulation (*j, i*) *∈ E* has a known monotonicity *l*(*j, i*) *∈* {⊕, ⊖}, but the input *j* of *F*_*i*_ does not exhibit this monotonicity; repaired by setting the monotonicity label to ? (unknown).

Note that in some instances, these issues may be intentional. For example, if an error of type 2 is detected such that the update function expression does not contain the incriminated variable at all, this could be a mechanism by which the authors specify that the relationship between the two variables is notable, but unused in this particular system. In such case, the *non-essential* annotation is the correct way to resolve this issue.

However, for the vast majority of errors we observed, this is not the case: The type 2 and 3 errors arose primarily from update functions that syntactically contain the problematic variable, but due to the structure of the function, either the variable does not contribute towards the output of the function at all, or the monotonicity of this contribution does not match the monotonicity declared by the authors.

### Unused components

Finally, we observe that many models admit variables that do not interact with the rest of the model in any meaningful way, or that such variables appear once all input essentiality errors are corrected. In general, we thus validate that the influence graph of the network in question is always weakly connected. That is, between any two variables, there must be some form of a direct or transitive relationship. If such disconnected components are found, we only retain the largest weakly connected component.

## Dataset Distribution

The dataset is managed through a versioned git repository that is continuously updated with new models. However, we regularly publish official *editions* of the dataset which freeze the set of models at a particular time-point and can be referenced separately.

### Model metadata

As already discussed, each model within a particular BBM edition is available in three formats, `bnet`^15^, `aeon`^16^, and `sbml`^17^, and with erased update functions on input variables. Furthermore, each model is accompanied by a collection of metadata which is provided both as a machine-friendly `json` file and a human-readable `markdown` document. As part of this metadata, we track the publication where the model first appeared (as a BibTeX reference), the original source of the model file, description of any modifications made to the model during validation, as well as the set of relevant keywords.

Using these keywords, we track the databases in which the model appears (e.g. `cell-collective`, `ginsim`, etc.) or methods used to construct the model (e.g. `casq`^22^). We also use the keyword `repaired` to label models where some logical inconsistencies were found during validation (the specific nature of the repair performed is detailed as a separate note), and keyword `multi-valued` to indicate that the model is a Booleanised version of a multi-valued network.

### Custom datasets

We recognise that our chosen model representation may not be suitable for all benchmarking and validation tasks. As such, we also include an interactive workflow which allows anyone to export a custom edition based on the BBM dataset. Specifically, this workflow allows the user to:

- Chose between the three model formats;
- Chose the desired representation of input variables (constant *true*, constant *false*, identity function, erased function);
- Filter the results using a custom Python expression evaluated for each model and its metadata.

Using this filter functionality, users can pick models based on structural properties (presence of a specific variable or regulation, the number of variables, connectivity, etc.), but also based on model metadata (model authors, keywords, etc.). To ensure reproducibility, each such custom dataset edition incorporates an additional metadata file with all parameters of the workflow, meaning the custom edition can be fully recreated based on this metadata file.

## Data Records

There are currently two editions of the BBM dataset, both available as Zenodo archives: edition 2021^23^ at the address https://doi.org/10.5281/zenodo.8020272, and edition 2022^24^ at https://doi.org/10.5281/zenodo.8020309. Any future updates to the dataset will be announced using the git repository at https://github.com/sybila/biodivine-boolean-models. Both editions contain the raw input model files, the generated output models and their metadata, as well as any programs necessary to validate the integrity of the models and to generate custom model editions. The models available in the latest edition (2022) are summarised in the Online-only Table 1.

## Technical Validation

First, to ensure compatibility of model formats, we validate the syntax of each model file using the tool `aeon.py`^21^ (some of the input model files even contain minor syntactic errors that had to be corrected). Furthermore, we already described the individual validation steps performed within our pipeline as part of the *Methods* section. This validation pipeline is executed for each commit of the aforementioned git repository, meaning no errors can be introduced into the dataset undetected.

However, we would like to emphasise the importance of this validation step by analysing the number and types of potential problems uncovered using our validation pipeline. This is summarised in Figure 2, which shows the number of problems grouped by type (with a more detailed breakdown in Online-only Table 1). Here, the *Other* category mainly covers models with syntactic issues or other obvious typos. What is especially alarming for us is the high number of regulations with invalid monotonicity or essentiality properties. Overall, we found that *∼* 1.5% of all regulations have potential integrity problems when compared to the actual update function.

**Figure 2.**
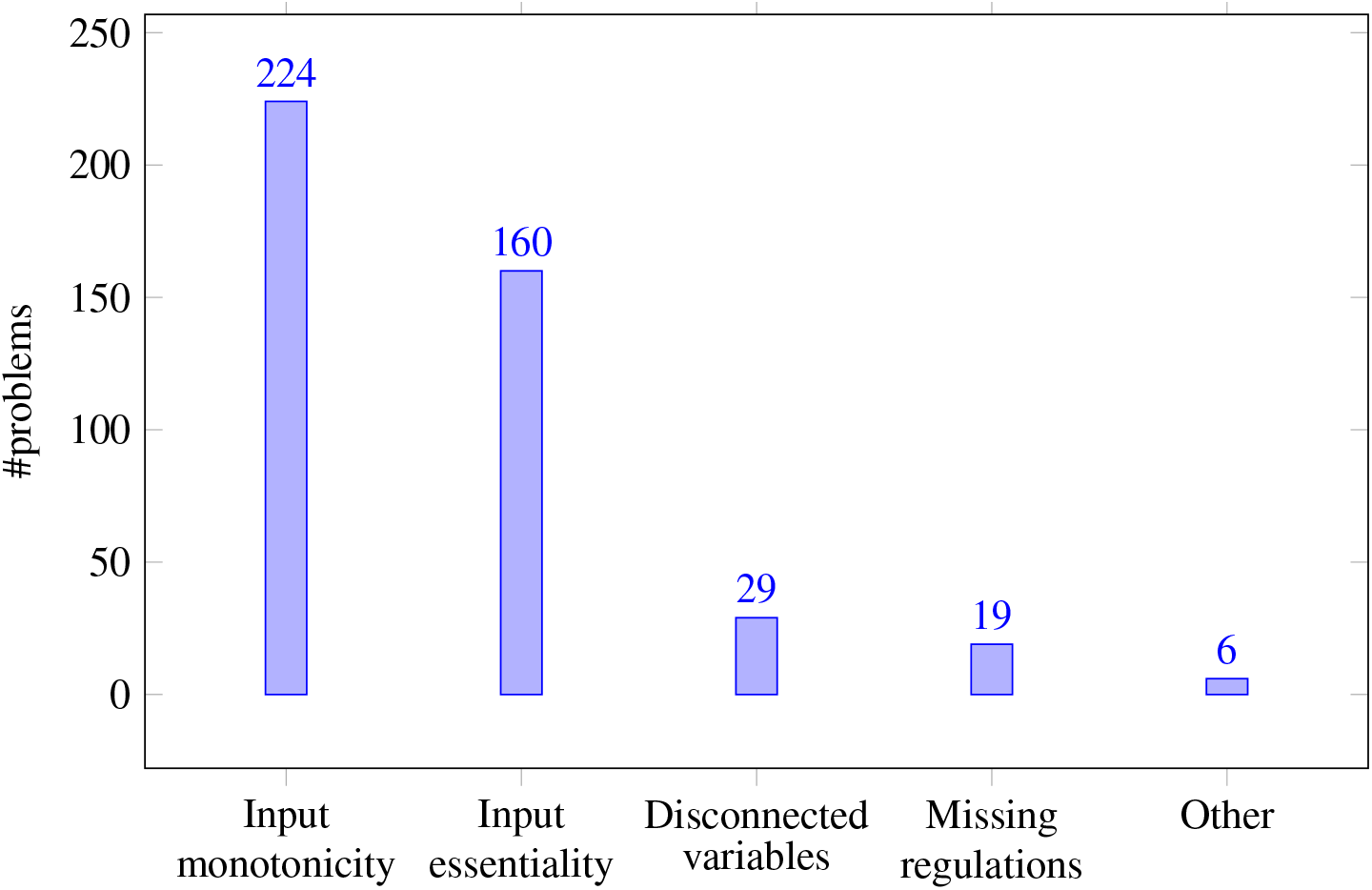
Types and multiplicity of individual problems uncovered by the validation step of the BBM pipeline (see Online-only Table 1 for per-model summary).

Finally, we should note that our validation still has potential blind spots and the total number of problems thus may not be final. First, the Booleanization process using `biolqm`^19^ which is applied to multi-valued networks always “resets” the network’s influence graph such that it is consistent with the Booleanized update functions. Since we currently cannot work with multi-valued networks directly, this means the two most prominent types of errors cannot be detected in these models. Second, many models do not come annotated with a signed influence graph and are only given as a list of functions. For such models, we thus have nothing to validate against. Nevertheless, we could still compare the employed regulations against other models and biological databases where applicable. Our goal is to address these shortcomings as part of the future work.

## Usage Notes

The model files distributed within BBM are compatible with many common tools for Boolean network analysis. In particular, this includes (but is not limited to) the tools available as part of the CoLoMoTo Jupyter notebook environment.^25^ Specifically, one can use the following tools (in the alphabetical order) to further analyse the properties of the published models:

- `aeon`^16^ and `aeon.py`^21^ perform symbolic analysis of the asynchronous network dynamics (attractors, fixed-points, reachability) and bifurcation analysis with respect to network inputs and parameters;
- `biolqm`^19^ and `ginsim`^14^ facilitate analysis of multi-valued models and other miscellaneous methods (translation between additional model formats, simulation, perturbation analysis, etc.);
- `cabean`^26^ facilitates target control and reprogramming of the network behaviour;
- `maboss`^27^ supports analysis of BNs under *stochastic* update semantics;
- `mpbn`^28^ computes the attractors and reachability properties in the *most-permissive* BN semantics;
- `pyboolnet`^15^ supports generation of random networks and synchronous simulation;
- `pystablemotifs`^29^ uses succession diagrams for attractor computation and network reprogramming;
- `trappist`^8^ computes the minimal and maximal trap spaces of the network as well as fixed-points.

For further reading, we recommend the repository of the CoLoMoTo notebook environment, which contains usage examples for most of the aforementioned tools distributed as user-friendly Jupyter notebooks.

## Code availability

The code, including previous revisions and all relevant model source files, is available as a git repository at the URL https://github.com/sybila/biodivine-boolean-models. Historical revisions of the source code associated with each BBM edition are also part of the aforementioned Zenodo packages. Submissions of new models are encouraged through issues and pull requests in this git repository (see instructions in the `CONTRIBUTING.md` file). The automated validation and post-processing relies on the tool `aeon.py`^21^, version `1.1.0`. Furthermore, `biolqm`^19^ was manually used for some of the initial model conversions, but is not used by the automated validation or the export pipeline.

## Supporting information

Online-only Table 2

Online-only Table 1

## Acknowledgements

This work was partially supported by GACR [grant No. GA22-10845S]; and Grant Agency of Masaryk University [grant No. MUNI/G/1771/2020]. This project has received funding from the European Union’s Horizon 2020 research and innovation programme under the Marie Skłodowska-Curie Grant Agreement No. 101034413.

## Figures & Tables

**Online-only Table 1**. List of all BBM models, including model ID, name, list of sources, and the corresponding publication. Also contains basic metadata of each model, such as the variable and regulation count.

**Online-only Table 2**. Associates BBM models with database entries. Lists a relevant BBM model reference for each entry in one of the tracked databases. Includes the date at which the model was last updated (in case of versioned database entries).

